# Revealing Early Spatial Patterns of Cellular Responsivity in Fiber-Reinforced Microenvironments

**DOI:** 10.1101/2024.01.12.575366

**Authors:** Saitheja A. Pucha, Maddie Hasson, Hanna Solomon, Gail E. McColgan, Jennifer L. Robinson, Sebastián L. Vega, Jay Milan Patel

**Affiliations:** Department of Orthopaedics, Emory University School of Medicine, Atlanta, GA, USA; Atlanta VA Medical Center, Department of Veterans Affairs, Decatur, GA, USA; Department of Orthopaedics and Sports Medicine, University of Washington, Seattle, WA, USA; Department of Mechanical Engineering, University of Washington, Seattle, WA, USA; Department of Biomedical Engineering, Rowan University, Glassboro, NJ, USA; Department of Orthopaedic Surgery, Cooper Medical School of Rowan University, Camden, NJ, USA

**Keywords:** Cell Sensing, Fiber-Reinforcement, Mechanobiology, Scaffold Microenvironment, Cell Heterogeneity

## Abstract

Fiber-reinforcement approaches have been utilized to replace aligned tissues with engineered constructs after injury or surgical resection, strengthening soft biomaterial scaffolds and replicating anisotropic, load-bearing properties. However, most studies focus on the macroscale aspects of these scaffolds, rarely considering the cell-biomaterial interactions that govern remodeling and ECM organization towards aligned neo-tissues. Since initial cell-biomaterial responses within fiber-reinforced microenvironments likely influence long-term efficacy of repair and regeneration strategies, here we elucidate roles of spatial orientation, substrate stiffness, and matrix remodeling on early cell-fiber interactions. Bovine mesenchymal stromal cells (MSCs) were cultured in soft fibrin gels reinforced with a stiff 100 µm polyglycolide-co-caprolactone fiber. Gel stiffness and remodeling capacity were modulated by fibrinogen concentration and aprotinin treatment, respectively. MSCs were imaged at 3 days and evaluated for morphology, mechanoresponsiveness (nuclear YAP localization), and spatial features including distance and angle deviation from fiber. Within these constructs, morphological conformity decreased as a function of distance from fiber. However, these correlations were weak (R^2^ = 0.01043 for conformity and R^2^ = 0.05542 for nuclear YAP localization), illustrating cellular heterogeneity within fiber-enforced microenvironments. To better assess cell-fiber interactions, we applied machine-learning strategies to our heterogeneous dataset of cell shape and mechanoresponsive parameters. Principal component analysis (PCA) was used to project 23 input parameters (not including distance) onto 5 principal components (PCs), followed by Agglomerative Hierarchical Clustering (AHC) to classify cells into 3 groups. These clusters exhibited distinct levels of morpho-mechanoresponse (combination of morphological conformity and YAP signaling) and were classified as High Response (HR), Medium Response (MR), and Low Response (LR) clusters. Cluster distribution varied spatially, with most cells (61%) closest to the fiber (0 – 75 µm) belonging to the HR cluster, and most cells (55%) furthest from the fiber (225 – 300 µm) belonging to the LR cluster. Modulation of gel stiffness and fibrin remodeling showed differential effects for HR cells, with stiffness influencing the level of mechanoresponse, and remodeling capacity influencing the location of responding cells. Overall, clustering of individual cells in stiff-soft microenvironments revealed spatial trends in cellular responsivity not seen by evaluating individual cell parameters as a distance from fiber alone.

**Impact Statement:** This study used PCA-AHC based clustering to identify MSC sub-groups from a heterogeneous population with distinct responses to stiff-soft microenvironments. Cell responsivity within a soft, fiber-reinforced fibrin gel microenvironment was influenced by the spatial localization of individual cells around a stiffer polyglycolide-co-caprolactone fiber. Additionally, modulation of gel substrate stiffness and matrix remodeling capacity further influenced the level of responsiveness and localization of responsive cell clusters around the fiber, which may contribute to scaffold design at the cellular level and foreshadow longer-term aligned tissue deposition.

## Introduction

Fiber-reinforcement for tissue engineering applications has grown substantially in the past decade^1^, with several advances in recapitulating the mechanical properties of load-bearing musculoskeletal tissues, such as the meniscus^2–5^, tendons^6^, and articular cartilage^7^. Generally, these strategies aim to mimic native aligned tissue by reinforcing a soft biological substrate with a stiffer polymer fiber network, typically composed of synthetic materials such as poly(lactic acid)^8^ or poly(ε-caprolactone)^9^. The softer substrate is often utilized for encapsulation of cells within natural materials such as collagen^8,10^, gelatin^11^, and fibrin^12^. Beyond aligned tissues, 3D printed scaffolds have gained popularity recently, especially as customizable, personalized implants^4,13,14^. While the field is moving closer to recapitulating the bulk mechanical properties of native tissues, the balance between biomechanical properties and tissue deposition at the cellular level remains challenging^1^. This is especially important to cell-laden fiber-reinforced gel scaffolds, where neo-tissue eventually bears load as the polymer fiber and substrate network degrades^2,3,5,15^ Many tissue engineering approaches involve fabrication and biomechanical testing of fiber-enforced scaffolds at the macroscale, yet they rarely consider the microscale cell-biomaterial interactions that likely govern eventual tissue formation. In fact, early dynamics of cell response, in terms of morphological characteristics and marker expression, have been known to mediate eventual tissue deposition and organization in aligned tissues, like the meniscus^16–18^. Furthermore, during development, the properties of the cellular microenvironment in aligned tissues are crucial in driving and maintaining cell phenotype^19–24^, and evidence suggests that early cell patterning can mediate extracellular matrix (ECM) architecture^16,25^ and ensuing tissue deposition. Therefore, early cell-biomaterial interactions and patterns of cellular mechanoresponse within these fiber-reinforced microenvironments require a deeper understanding that would aid in optimization of fiber-reinforcement strategies for the repair and replacement of aligned musculoskeletal tissues^14^.

Although morphological and mechano-responsive patterns in 3D matrix microenvironments have been well studied, there remains a knowledge gap in the nuances of 3D matrix mechanosensing^26–28^. Understanding cellular matrix mechanosensing in 3D microenvironments is critical to gaining insight into factors governing cell behavior *in vivo*, especially in the context of musculoskeletal load-bearing tissues. Towards this, several studies have used ECM-mimetic hydrogel systems, such as fibrin and hyaluronic acid^29–31^. Fibrin, in particular, mimics early wound healing environments that are readily remodeled, exhibits excellent biocompatibility, and is finely tunable^28,32^. Thus, in this study, we utilized fibrin as the soft biological substrate for cells to remodel and respond within a fiber-reinforced gel. Response to and remodeling of the surrounding matrix can be influenced by several factors, leading to variable patterning and differentiation within the microenvironment^16,22,25,33^. Understanding the dynamics by which cells sense within an anisotropic 3D environment can inform design strategies for fiber-reinforced scaffolds at the microscale. Specifically, cells can sense and respond to microenvironmental cues, such as anisotropic structural orientation and depth^34^, biophysical properties of the surrounding matrix (e.g., degradation and mechanics)^15,35^, and proximity to rigid materials (e.g., polymer fiber) in stiff-soft environments^36^. Through mechanosensing of these cues, downstream morphological behavior and patterning of the fiber-reinforced microenvironment can govern eventual tissue deposition and architecture^37,38^. The transcriptional regulator YAP (Yes-associated Protein) serves as a mediator of matrix mechanosensing^27,39,40^, correlating to dynamics of ECM deposition^27,39–42^. Therefore, by measuring early YAP nuclear localization and cell morphological features, we can reveal holistic patterns of cell response in stiff-soft fiber-reinforced microenvironments.

Perhaps the most widely utilized cells in musculoskeletal tissue engineering are mesenchymal stromal cells (MSCs), but heterogeneity amongst these cell populations must be acknowledged to study their therapeutic potential^43–45^. Because of this, organizing heterogeneous populations of MSCs based on their functional potential is crucial to accurately evaluating treatments on a single-cell level. Several studies have begun to utilize machine-learning based modeling techniques to group cell populations by response^41^, morphology^46^, or marker expression^45^, providing insight into the heterogeneity of MSCs within cell-biomaterial microenvironments. Through these techniques, cell populations of interest can be highlighted to better reveal response patterns that are influenced by interactions at and near the cell-biomaterial interface.

The goals of this study are to 1) investigate spatial patterns of cell response in a fiber-reinforced microenvironment; 2) utilize a machine-learning based clustering approach to identify spatially responsive sub-groups within a heterogeneous cell population; and 3) understand how alterations of this microenvironment mediate these patterns of response. Together, these experimental results displayed the applicability of machine learning in revealing patterns in a heterogeneous environment and the importance of spatial orientation, substrate biophysical properties, and matrix remodeling in governing early cell response in fiber-reinforced microenvironments.

## Materials and Methods

### Cell Preparation and Culture

Marrow stromal cells (MSCs) were isolated from a juvenile (1–3 weeks old) bovine stifle joint (Research 87, Boylston MA). Briefly, subchondral trabecular bone blocks were shaken in heparin-containing (0.2% w/v heparin) Dulbecco’s modified Eagle’s medium (DMEM), and the resulting solution was centrifuged, resuspended in basal medium with 10% fetal bovine serum (FBS; VWR 97068-085; Lot: 274K20), and plated for expansion. Cells were then frozen in liquid nitrogen, thawed, and expanded as needed. Passage 1 cells that were only expanded once after thawing were used for all experiments.

### Fibrin gel fabrication and mechanical testing

Prior to cell experiments, acellular fibrin gels were fabricated to evaluate mechanical properties. Fibrinogen from bovine plasma (Sigma F8630; 10, 25, and 50 mg/mL final concentration) and thrombin from bovine plasma (Sigma T4648; 5 U/mL final concentration) were mixed with 200 mM calcium chloride (CaCl_2_) in phosphate-buffered saline (PBS) to fabricate gels. To do so, a mixture of thrombin/CaCl_2_/PBS (5 µL) was first pipetted onto a glass slide. Next, 5 µL of fibrinogen was pipetted directly onto this drop, pipetting several times to mix prior to fibrinogenesis. Gels were incubated for 1 hour at 37 °C and 5% CO_2_ and suspended in 1X PBS. Gel mechanics were characterized via nano-indentation (Optics 11 Pavone) with a 27 µm-radius spherical probe (0.020 N/m). At least 20 points were tested on each gel, and load-indentation curves were fit with a Hertzian model to obtain an effective Young’s modulus.

### Cell encapsulation and hydrogel culture

To encapsulate MSCs within fibrin gels, passage 1 MSCs were thawed and expanded in basal medium for 5-7 days, until ∼75% confluent in 100 mm cell culture dishes. Cells were then trypsinized and added into the thrombin/CaCl_2_/PBS solution, reaching a final concentration of 1.25 million cells/mL (12,500 cells per 10 µL gel). Gels composed of varying concentrations of fibrinogen (10, 25, or 50 mg/mL) were prepared in an 8-well chambered glass slide (**Fig. 1A**), followed by addition of a 5-0 MonoQ fiber (polyglycolide-co-caprolactone; [PGCL]; ∼100 µm diameter; Ethicon) directly into each droplet prior to gelation (1 hour, 37 °C, 5% CO_2_). Following incubation, gels were cultured in chemically defined medium (DMEM, 1% v/v penicillin-streptomycin-fungizone [PSF], 50 µg/mL vitamin C, 0.1 mM dexamethasone, 40 µg/mL L-proline, 100 µg/mL sodium pyruvate, 0.1% v/v ITS liquid medium supplement, 1.25 mg/mL bovine serum albumin [BSA], 5.3 µg/mL linoleic acid [LA]) supplemented with 10 ng/mL TGF-β3 and low or high concentrations of aprotinin (10 or 100 KIU/mL). Aprotinin inhibits fibrinolysis by preventing plasmin from inducing degradation of crosslinked fibrin. Samples were cultured for three days (37 °C, 5% CO_2_), and medium was replaced after 2 days in culture.

**Figure 1.**
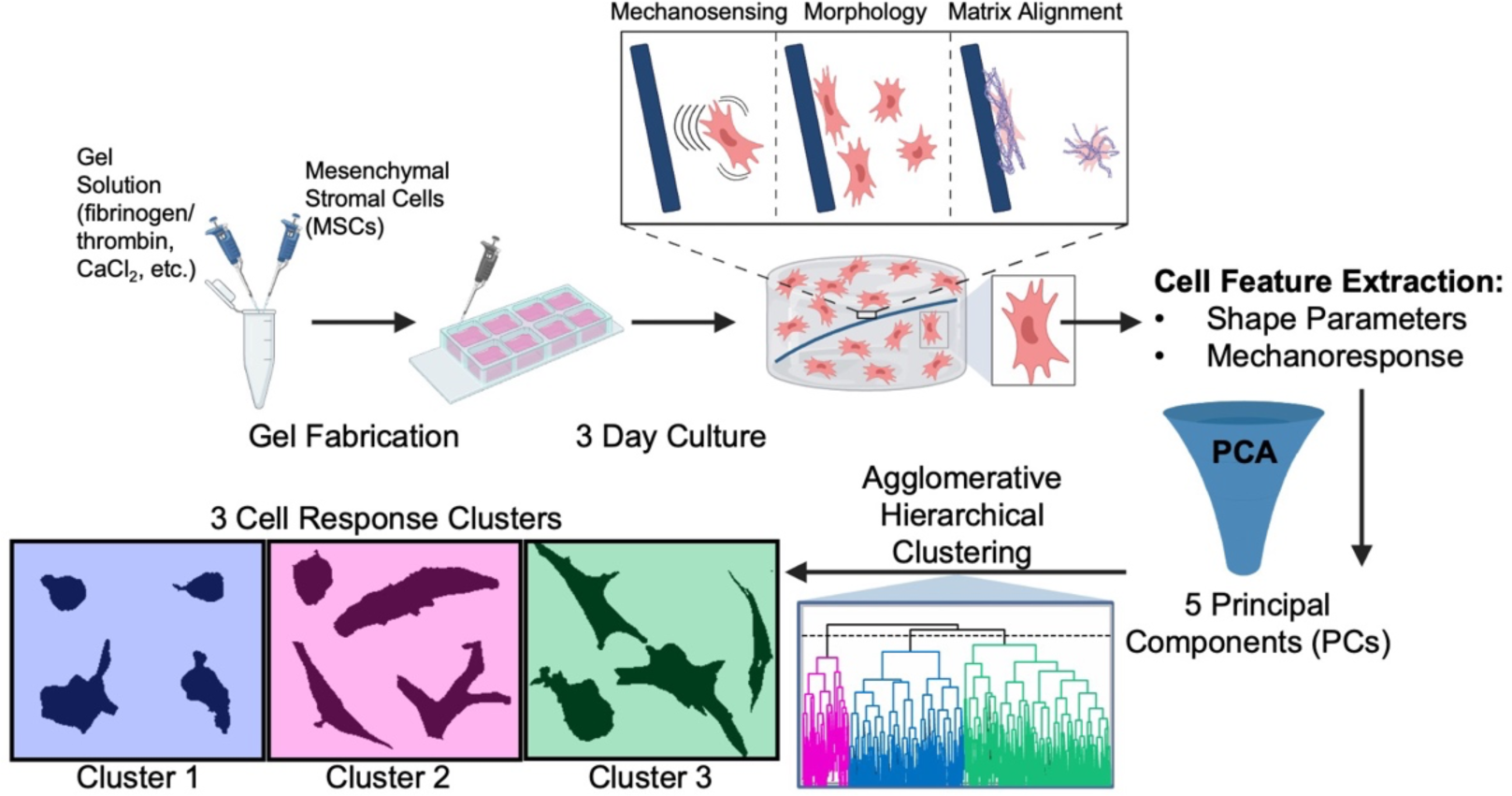
Experimental Design, cell feature acquisition. [A] Simulation of fiber-reinforced microenvironment via fibrin gel fabrication and culture. [B] Cell feature extraction (n=23) and Principal Component Analysis (PCA) to project onto 5 Principal Components (PCs). [C] Clustering of total cells (n=943) into three distinct clusters based on level of response. (Created with BioRender.com)

### Immunofluorescent Staining

Following culture, gels were rinsed in PBS and fixed in 10% Carson’s buffered formalin for one hour at room temperature (RT). Next, gels were rinsed thrice in PBS and permeabilized (1% Triton X-100 in PBS) for 45 minutes. After three PBS rinses, gels were blocked (3% BSA in PBS) for 30 minutes. Next, gels were stained for YAP1 (Invitrogen PA5-87568; rabbit polyclonal; 1:200 in 1% BSA) for 60 minutes at RT. Following three PBS rinses, gels were stained with secondary antibody (AlexaFluor 647, goat anti-rabbit; 1:200) and Phalloidin (AlexaFluor 555, 1:400) in 1% BSA for 60 minutes at RT. Gels were rinsed again and nuclei were stained with DAPI (1:1000 in 1% BSA) for 20 minutes at room temperature. Chambers were removed, and samples were mounted with Prolong Gold and a rectangular coverslip (24 mm x 50 mm) was placed on top.

### Imaging and Cell Parameter Extraction

A Nikon A1R confocal microscope was used to visualize cells in fiber-reinforced gels. Cells within 300 µm of the fiber were used for analysis. ND2 images were first processed in ImageJ FIJI to obtain maximum intensity projections and split channels (DAPI, phalloidin, YAP) into individual TIFF files. Next, a custom CellProfiler pipeline was created to identify individual nuclei (from the DAPI channel), their encompassing cells (from the phalloidin channel), and single-cell YAP intensities in the nucleus, cytoplasm, and perinuclear ring using the YAP channel. From these images, 23 measurements were acquired for each identified cell (n = 943 cells). Cell and nuclear shape measurements included: Area, Compactness, Eccentricity, Form Factor (4*π*Area/Perimeter^2^), Major Axis Length, Max Feret Diameter, Minor Axis Length, Perimeter, Solidity, Angle Deviation, and Aspect Ratio. YAP parameters measured were nuclear/cytoplasmic intensity ratio and nuclear/perinuclear intensity ratio. In addition, distance from each cell nuclear centroid to its closest point on the polymer fiber were acquired for each measured cell. From these raw cell features, conformity index (Eq.1) and morpho-mechanoresponse (Eq.2) parameters were calculated for each cell. Conformity index describes the degree to which individual cells align to the morphology of the stiff polymer fiber, considering both cell spreading and cell angle to output a value between 0 (less conforming) and 1 (more conforming). Morpho-mechanoresponse considers both conformity index and YAP nuclear/cytoplasmic intensity ratio to output a singular value describing the degree to which individual cells sense and conform to the stiff polymer fiber.

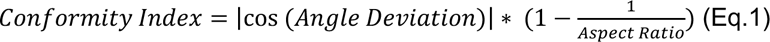

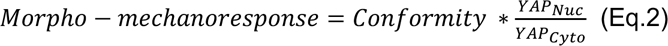

### Analysis Pipeline, PCA, and AHC

After acquisition of 23 parameters for 943 cells in one of six fiber-reinforced gel groups (25, 50, or 100 mg/mL fibrinogen in medium supplemented with 10 or 100 KIU/mL aprotinin), data were standardized by subtracting the mean from each value and dividing by standard deviation. Next, PCA using a correlation matrix was run on this set of 943 cells using XLSTAT software (Addinsoft Corporation), outputting five principal components (PCs; **Fig. 1B**), comprising 83.78% of the variability present in the total data set (80% variability was used as a threshold). Weighted factor scores for each PC were outputted for each individual cell. Next, the five PC scores for each cell were used for Agglomerative Hierarchical Clustering (AHC) analysis in XLSTAT. Clusters were determined using dissimilarity (Euclidian Distance), and Ward’s method was used for agglomeration of clusters. Using the Hartigan Index, three was determined as the optimal number of clusters to ensure maximum distance between clusters^47,48^. AHC divided the heterogeneous set of 943 cells into cluster 1 (n = 351), cluster 2 (n = 450), and cluster 3 (n = 142). These clusters were used to distinguish patterns of cell response in different cell families (**Fig. 1C**).

### Statistics

Linear regression analysis was used to analyze scatterplot data. For column graphs, outliers were identified using the ROUT method, and normality was tested with the D’Agostino & Pearson test. Next, the Kruskal-Wallis non-parametric ANOVA test (distributions were found to be non-normal) was used to compare groups with 2 or more comparisons, using Dunn’s multiple comparisons test to determine significance between groups. The Mann-Whitney non-parametric t-test was utilized for comparisons between two groups. In all violin plots, distribution of individual data points is shown, the bold middle line in each plot indicating median, and two dotted lines above and below indicating quartiles. In each graph, *, **, ***, and **** represent p < 0.05, 0.01, 0.001, and 0.0001, respectively.

## Results

### Relationships between cell features and fiber distance within stiff-soft microenvironments are highly heterogeneous

Following 3-days of MSC culture within fiber-reinforced microenvironments, spatial heterogeneity was present in terms of single-cell conformity and mechanoresponsiveness. Considering several cell shape parameters such as angle deviation from fiber, aspect ratio, and conformity, considerable variability was seen on a cell-to-cell basis (**Fig. 2A**). Both conformity and YAP nuclear localization values for cells within the entire population (n = 943) exhibited a statistically significant negative correlation (p = 0.0017, p < 0.0001, respectively) with distance from the fiber (**Fig. 2B & 2C**). Consequently, morpho-mechanoresponse values for this set of cells also exhibited a negative correlation (**Fig. 2D**; p < 0.0001). However, all three of these correlations were weak (R^2^ < 0.1) despite a statistically significant deviation from a slope of 0, motivating the need for machine learning-based higher-order strategies to parse through heterogeneity within the data set.

**Figure 2.**
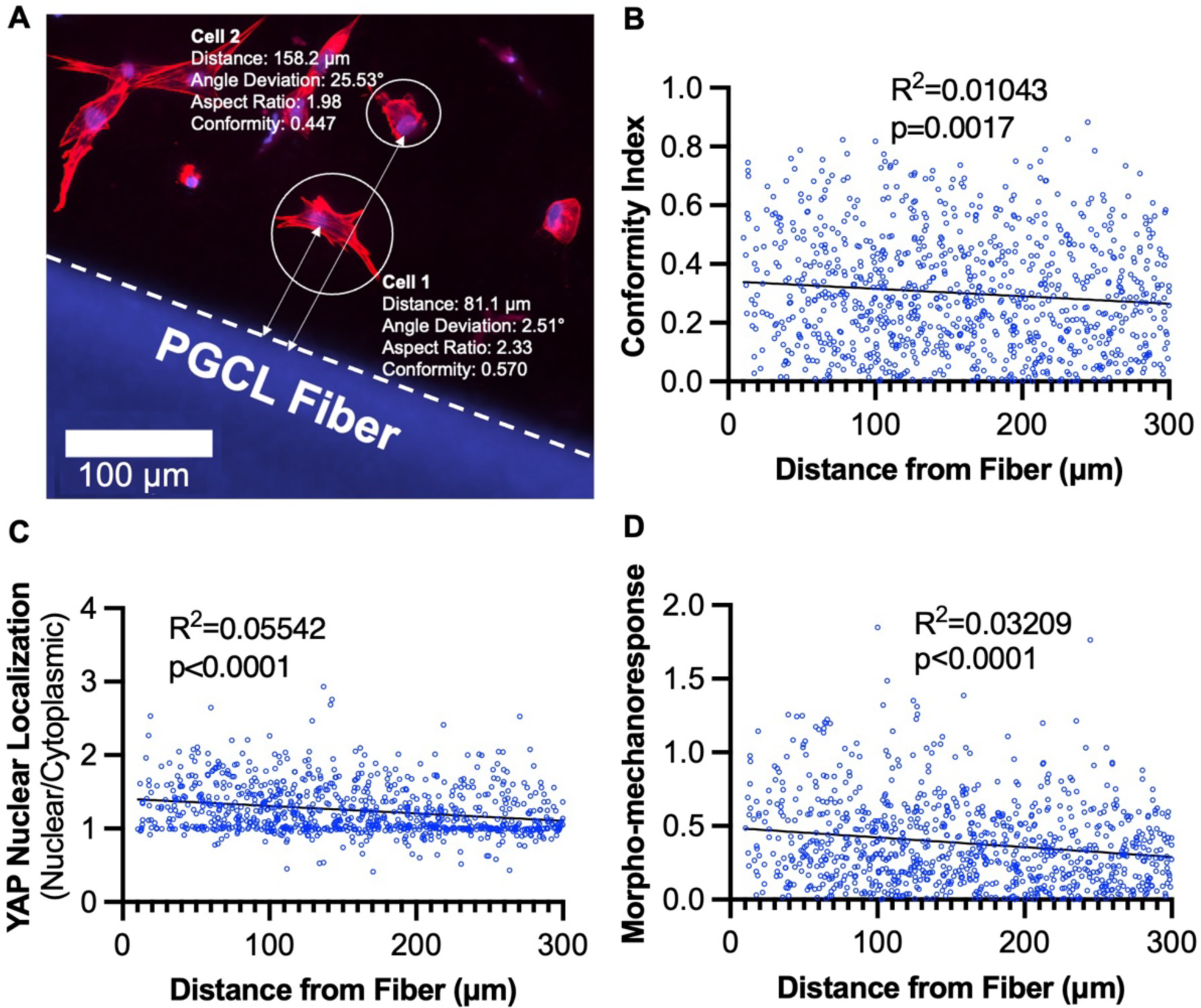
Cell shape and nuclear response to a stiff polymeric fiber in a fiber-reinforced environment is highly heterogeneous. [A] Distance, angle deviation, and YAP Nuclear/Cytoplasmic Ratio measurements of individual cells around a polymer fiber. [B] Conformity index, [C] YAP Nuclear Localization, and [D] Morpho-mechanoresponse values of cells (n=943) within a 300 µm area around a fiber as a function of distance from the fiber.

### 3D cell mechanoresponse decreases in a stiffness-dependent manner

To investigate the influence of gel substrate mechanics on patterns of cell behavior in the fiber-reinforced microenvironment, MSCs in stiff-soft environments comprised of varying fibrinogen concentrations (50, 25, and 10 mg/mL) and low aprotinin (10 KIU/mL) were evaluated. As expected, lower fibrinogen concentration resulted in lower gel stiffness (**Fig. 3A**). MSCs in lower stiffness (10 mg/mL fibrinogen) gels also exhibited greater morphological and nuclear response, with slightly higher conformity index values and significantly higher YAP nuclear localization (**Fig. 3B & 3C**). Overall, morpho-mechanoresponse was significantly greater for cells encapsulated in 10 mg/mL fibrinogen fibrin gels (**Fig. 3D**), indicating the importance of substrate mechanics in a cell’s ability to respond to biophysical stimuli at the cell-fiber interface.

**Figure 3.**
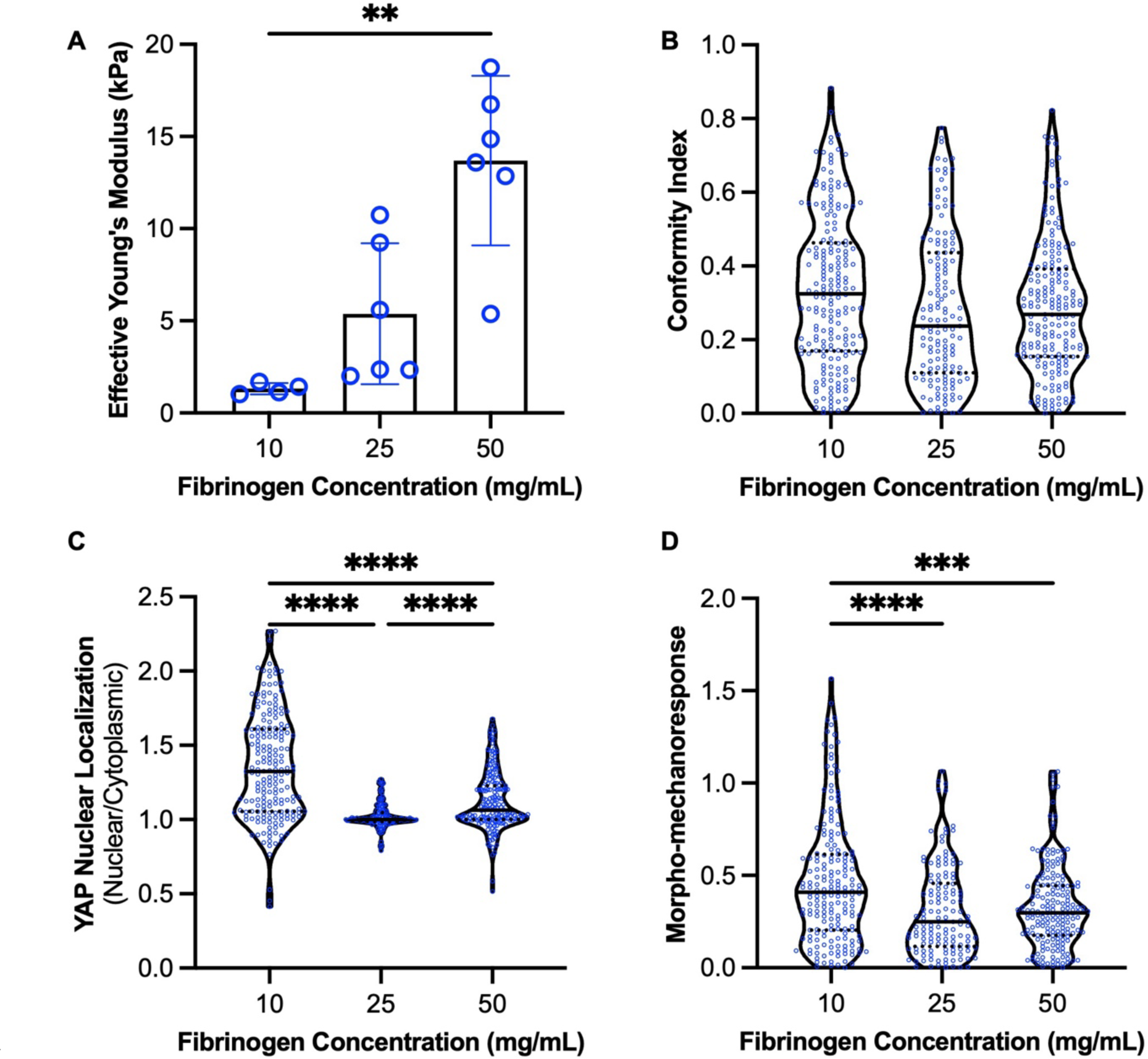
Cell response to polymeric fiber is heightened in lower-stiffness (lower fibrinogen content) fibrin constructs. [A] Effective Young’s modulus of gels fabricated with varying fibrinogen concentrations. [B] Conformity index of cells in varying Fibrinogen Concentration fibrin gels (all low aprotinin) [C] YAP nuclear localization ratio of cells in varying Fibrinogen Concentration fibrin gels (all low aprotinin) [D] Morpho-Mechanoresponse of cells in varying Fibrinogen Concentration fibrin gels (all low aprotinin) **, ***, **** represent p<0.01, 0.001, 0.0001, respectively.

### PCA-AHC analysis is able to separate heterogeneous MSC populations into three sub-groups

Due to the high heterogeneity observed in the holistic analysis of our large MSC population, we employed a PCA-AHC clustering approach to separate the cells into distinct sub-groups. We first projected 23 input variables calculated for each cell to 5 principal components (PCs) which comprised 83.78% of the cumulative variability in the data set (**Fig. S1**). These input variables each contributed differentially to each PC (**Fig. 4A**), with each PC correlating to each parameter to varying extents (**Fig. 4B**). PC1, responsible for 37.54% of the variability, was relatively evenly contributed by all parameters (largest contributor: major axis length, 8.17%), with cell and nuclear length parameters (axis length, perimeter, area) contributing most significantly. PC2, responsible for 20.60% of the variability, similarly correlated to greater cell length parameters, but correlated negatively to nuclear length parameters. Interestingly, PC4 (7.15% variability) was strongly linked to YAP parameters (∼40% contribution each), and PC5 (6.31% variability) was strongly linked to cell and nuclear angle deviation (∼46% each). Together, these PCs were used for agglomerative hierarchical clustering into three distinct clusters, shown by a PC1 versus PC2 plot in **Fig. 4C**. These three clusters also demonstrated mild separation when PC1 was plotted versus distance from fiber (**Fig. 4D**).

**Figure 4.**
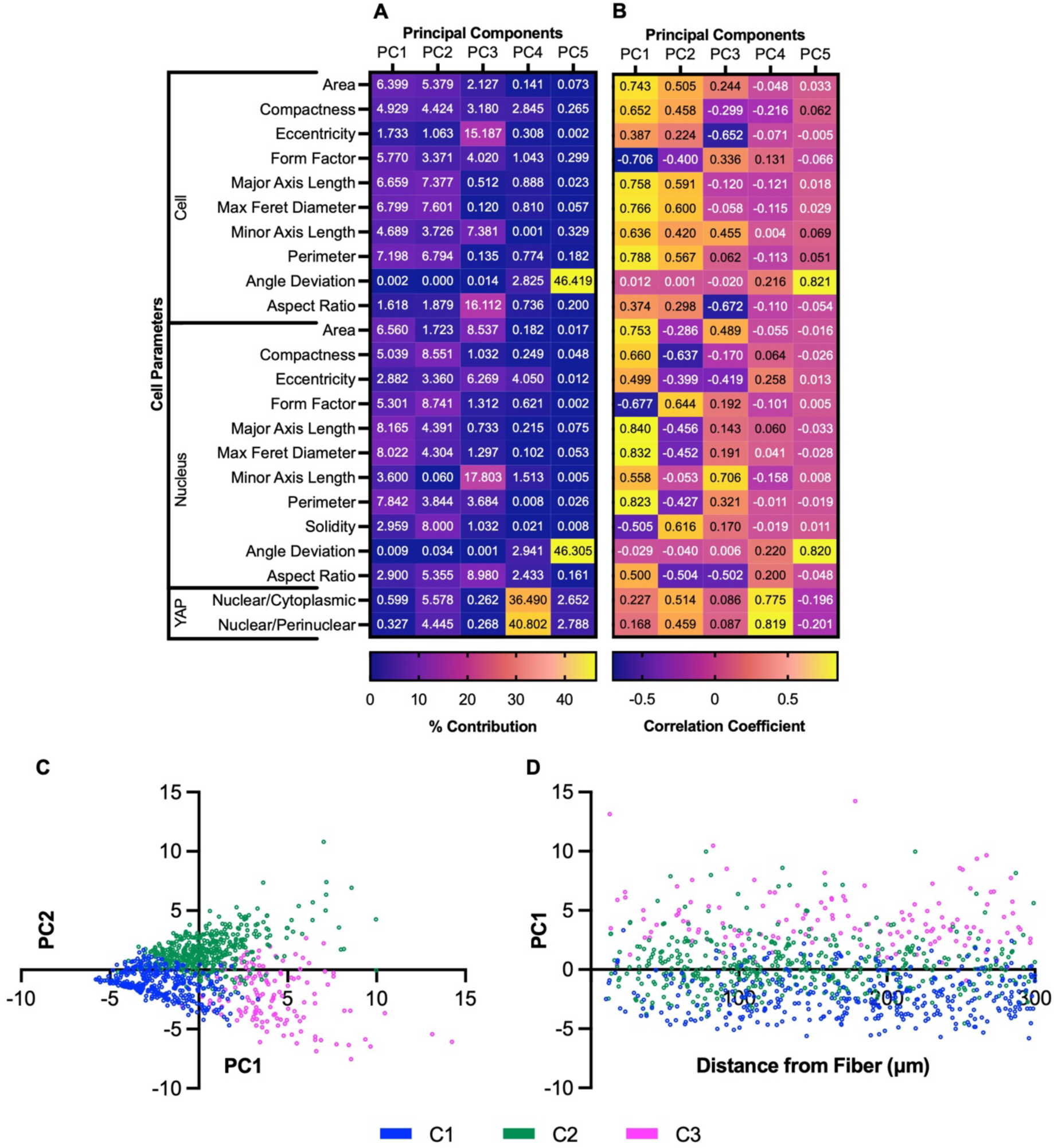
A heterogeneous set of cells within a simulated fiber-reinforced hydrogel microenvironment can be clustered using Principal Component Analysis followed by Agglomerative Hierarchical Clustering. [A] Contributions of each cell parameter to each principal component by percentage. [B] Correlations between each cell parameter and principal component. [C] Separate Cell clusters depicted on a PC2 vs. PC1 factor score plot. [D] PC1 factor score of cells from 0-300 µm from fiber, colored by cluster.

### PCA-AHC analysis reveals MSC clusters with distinct cellular features

The three clusters identified from the PCA-AHC analysis were compared with regards to morphological and mechanoresponse metrics. Cluster 2 cells displayed the highest conformity index measurements, followed by cluster 3, then cluster 1 (**Fig. 5A**). Cluster 2 cells also displayed the highest levels of YAP nuclear localization (p < 0.0001), and cluster 1 cells exhibited the lowest levels centered around a YAP nuclear ratio of 1. Cluster 3 cells exhibited a YAP response between clusters 1 and 3 (**Fig. 5B**). Considering both morphological and mechanosensing responses, cluster 2 cells exhibited the greatest morpho-mechanoresponse followed by cluster 3 cells and cluster 1 cells, with all groups demonstrating statistically significant differences from one another (**Fig. 5C**). A cumulative distribution plot of morpho-mechanoresponse measurements further highlights this separation between clusters (**Fig. 5D**). Based on these findings, the clusters were renamed by increasing morpho-mechanoresponse (Cluster 1 labeled LR for Low Response; Cluster 2 labeled HR for High Response; Cluster 3 labeled MR for Medium Response) for subsequent spatial analyses.

**Figure 5.**
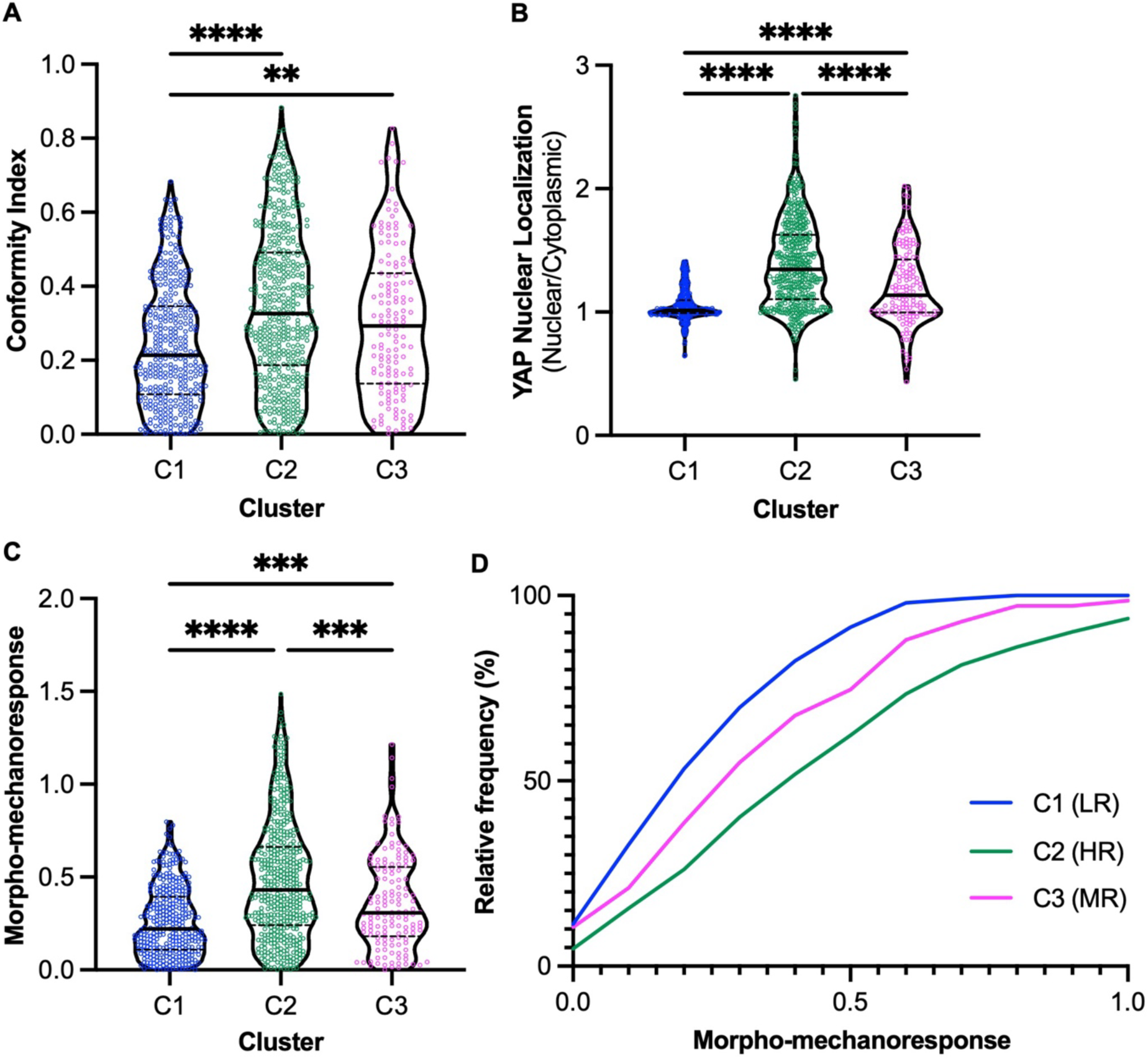
Classification of acquired cell clusters based on cytoskeleton and nuclear response parameters. HR=High Response, MR=Medium Response, LR=Low Response. [A] Conformity index of cells in each cluster (0-1) [B] YAP nuclear localization of cells in each cluster [C] Morpho-mechanoresponse of cells in each cluster. [D] Cumulative Distribution plot depicting the Morpho-mechanoresponse of cells in each cluster. Clusters assigned as: C1 = Low Response (LR), C2 = High Response (HR), C3 =Medium Response (MR). **, ***, **** represent p<0.01, 0.001, 0.0001, respectively.

### Cells localized to rigid fibers in stiff-soft microenvironments according to cluster responsiveness

After clustering the heterogeneous MSC data set into HR, MR, and LR groups, spatial patterning of these cell families around the fiber was examined. MSCs in the HR cluster appeared to localize closer to the fiber, while cells in the LR group seemed to localize further, suggesting a distance-dependent trend of response (**Fig. 6A**). Upon quantification, HR cells were located significantly closer to the fiber compared to MR and LR cells (**Fig. 6B**). In terms of distribution of cells in each cluster, the frequency distribution of HR cells was right skewed (skewness = 0.39), while the distributions of MR and LR cells were left skewed (skewness = -0.25, -0.24, respectively; **Fig. S2**), further illustrating the distance-dependent patterning of the microenvironment. Furthermore, when separating the 300 µm region of interest into four 75 µm bins, the proportion of HR cells showed a clear decrease from the first to fourth bin. In contrast, LR cells showed the opposite trend (increase from first to fourth bin), while MR cells seemed to be scattered evenly throughout the stiff-soft microenvironment (**Fig. 6C**).

**Figure 6.**
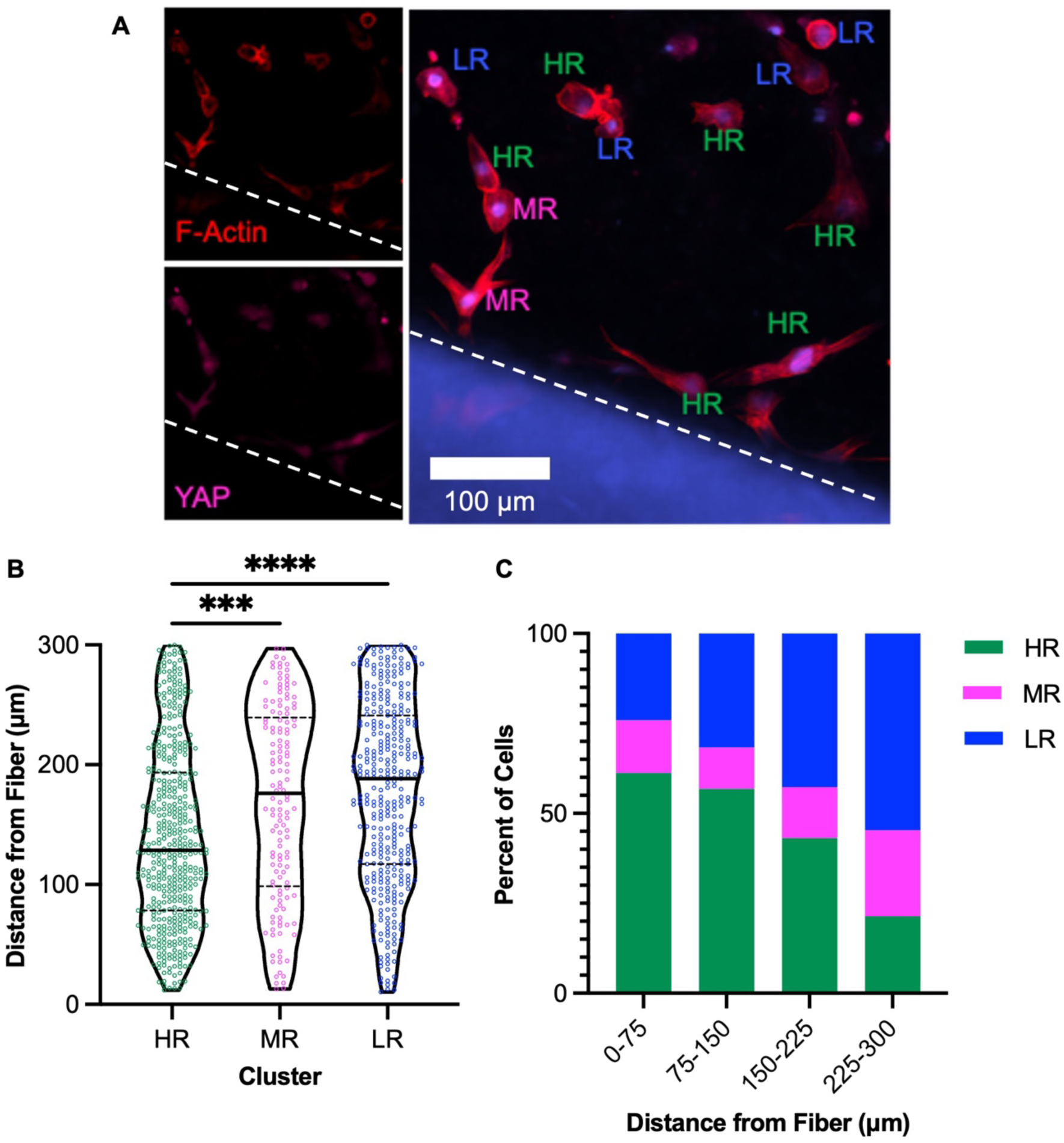
Cell clusters localize differentially in fiber-reinforced micro-environment. [A] Cells proximal to fiber labeled by cluster. HR=High Response, MR=Medium Response, LR-Low Response [B] Distances of all cells (n=943) from each cluster from the fiber. [C] Percentages of each cell cluster (All Cells) within different regions around fiber. *, ***, **** represent p<0.05, 0.001, 0.0001, respectively.

### Stiffness and matrix remodeling influence cell response and localization

After establishing that the MSCs in the HR, MR, and LR clusters localized to prescribed distances from the fiber in stiff-soft microenvironments, we next sought to use the identified clusters to better investigate the impacts of gel stiffness (controlled by fibrinogen concentration) and gel remodeling (modulated by aprotinin dosage) on cell responses. In HR cells specifically, morpho-mechanoresponse of cells was significantly increased in softer (lower fibrinogen concentration) gel environments (**Fig. 7A**). However, no significant differences were seen in terms of location of HR cells, relative to the fiber, in different stiffness environments (**Fig. 7B**). Similar trends, albeit not as statistically significant, were seen in MR and LR cells (**Figs. S3A, S3B**). Interestingly, higher aprotinin dosage over three days had no significant effect (p = 0.7081) on morpho-mechanoresponses of HR cells (**Fig. 7C**) but led to significantly (p = 0.0005) closer localization to the fiber (**Fig. 7D**). Again, similar trends were seen in other clusters in terms of morpho-mechanoresponse, but not distance (**Figs. S4A, S4B**).

**Figure 7.**
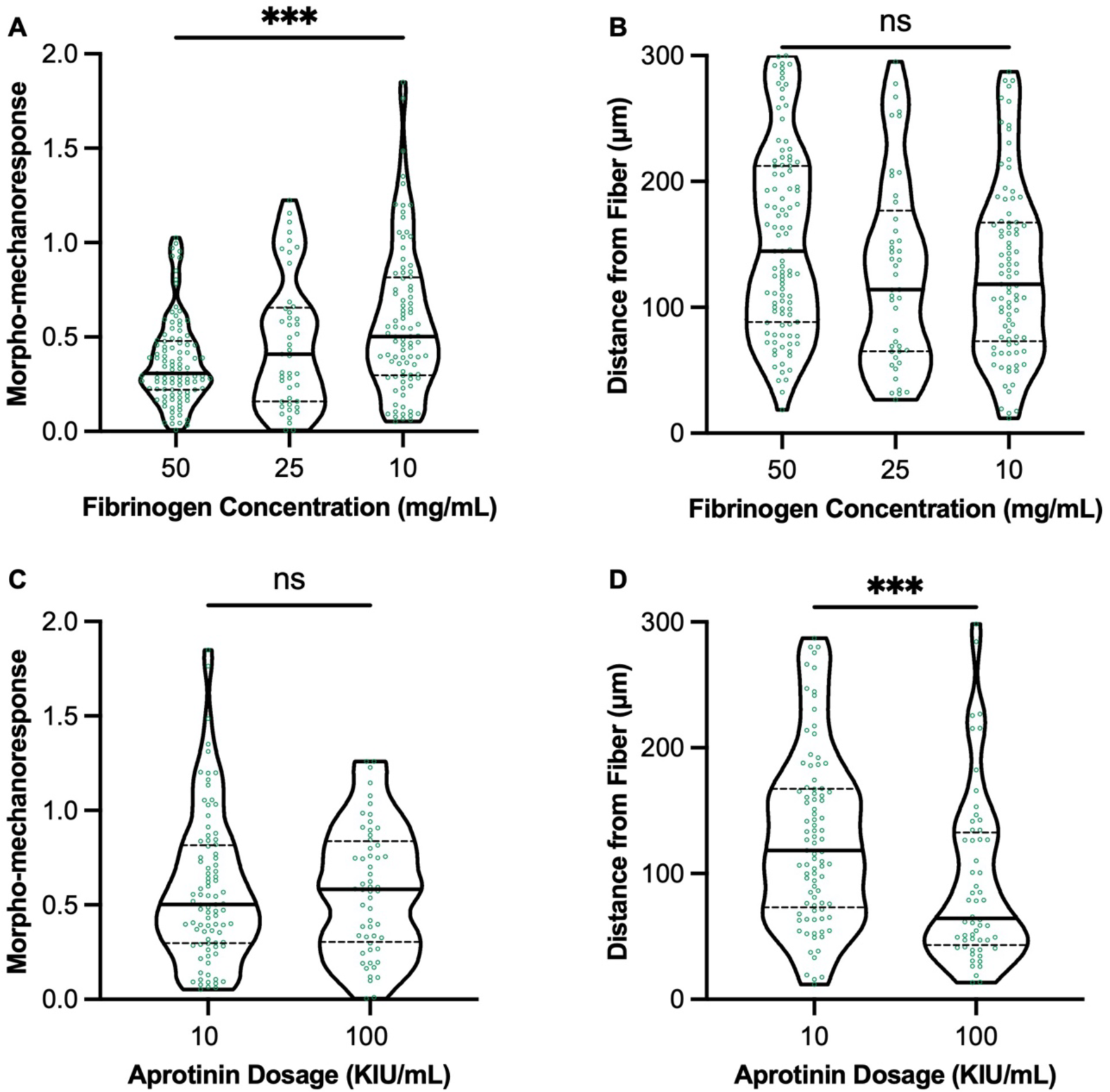
Stiffness of hydrogel substrate and fibrin remodeling capacity differentially influence level of morpho-mechanoresponse and spatial response of highly responsive cells in fiber-reinforced microenvironment. [A] Morpho-mechanoresponse and [B] Distance from fiber of HR cells in fibrin gels of varying fibrinogen concentration. [C] Morpho-mechanoresponse and [D] Distance from fiber of HR cells in fibrin gels supplemented with 10 and 100 KIU/ml concentrations of aprotinin. *** represents p<0.001.

## Discussion

The design and implementation of fiber-reinforced scaffolds to replace aligned tissues has gained popularity in recent years^1,4,6,8–11^, but the cell-scale patterns of responses within fiber-reinforced microenvironments is poorly understood. In this study, we investigated spatially dependent responses of single cells in these environments and the influence of substrate stiffness and degradation on morpho-mechanoresponses. Our cell clustering approach identified distinct MSC sub-populations within stiff-soft microenvironments, shed new light on spatial patterns of cell response, and increased our understanding of how the interplay between microenvironment stiffness and remodeling can tune cellular response patterns.

Heterogeneity in MSC populations has been well documented in the field^43–45^. Certainly, in terms of bone-marrow derived cells, cellular subsets are present, further owing to the heterogeneity present in the raw, unsorted marrow-derived preparations in this study. From spatial data of MSC morphological response (Conformity Index) and mechanoresponse (YAP), a distance dependent correlation was observed, with response decreasing with distance from the fiber. However, the weak correlations reduced confidence in generating conclusions about patterns of cellular responsivity within the fiber-reinforced microenvironment, motivating the need to use machine learning principles, as done in other single-cell response studies^41,45,46^. In the present study, PCA was performed to reduce 23 cell and nuclear parameters to a lower-dimension space. A cumulative variability threshold of 80% was set prior to analysis in order to preserve most of the variability of the data set while also reducing dimensionality, resulting in five principal components being used for clustering analysis (83.78% variability). An agglomerative hierarchical clustering approach successfully divided our heterogeneous cell population into three distinct clusters with varying levels of response (HR, MR, LR), based on conformity index and morpho-mechanoresponse. Through this clustering approach, the heterogeneous MSC population, composed of various uncharacterized cell types, as well as both senescent and non-senescent MSCs^45^, were divided to highlight key cell sub-groups of interest.

Upon organization of these cell clusters, we obtained an understanding of spatial patterns of cell localization in different cell groups. Hyde et al. demonstrated that cell patterning during meniscal tissue formation may mediate ECM organization^25^, contributing to the significance of spatial patterns seen in this study. The distance-dependent organization of more responsive cells dominating areas closer to the fiber provide insight into the spatial dynamics of the early fiber-reinforced microenvironment, which may contribute to the architecture of mature, aligned tissues^16,25^. The interplay between low and high response cells in terms of their localization within stiff-soft microenvironments can be crucial to optimizing fiber-reinforced technologies at the cellular level to maximize organization and alignment of eventual deposited tissue. Specifically, we have explored how the depth by which cells can sense a rigid fiber in a stiff-soft environment^34,36^ can influence spatial patterning of the 3D fiber-reinforced microenvironment. By illustrating the distance-dependent nature by which cells spatially sense rigid material across a soft medium, we have provided insight into the dynamics of cell instruction, in terms of orientation and differentiation, by topological features of the cell-biomaterial network. These dynamics can inform scaffold design at the microscale, from fiber spacing/diameter to methods for cell encapsulation to maximize sensing.

The physical properties of the fiber-reinforced microenvironment, namely substrate stiffness and 3D matrix remodeling, can have major implications on cell matrix mechanosensing and differentiation^9,22,24,26,28,33,49,50^. By modulating these factors in this study, we have shown novel evidence that, in the context of response to a stiff reinforcing polymer fiber, substrate stiffness predominantly mediates the extent to which responsive cells respond and conform (“how”), while the ability to remodel 3D matrix primarily influences the distance to which the cells can respond (“where”). The unique interplay between substrate stiffness and remodeling can be utilized to finely govern cell response in a fiber-reinforced microenvironment, controlling ultimate tissue deposition and architecture^15,16,22,23,25,35^. To optimize fiber-reinforced scaffold technology for anisotropic tissue regeneration^14,38^, the cellular nuances at the microscale and microenvironmental factors that govern these nuances must be considered^26,33^. Together, the patterns of cell response we have provided can be applied to microscale design of fiber-reinforced tissue engineering approaches optimizing early cell instruction to enhance proper tissue maturation at the macroscale through manipulation of substrate topography, fiber size/orientation, and matrix remodeling.

The results in this study are of interest to the tissue engineering field but present some limitations. For example, while the cellular heterogeneity within our simulated fiber-reinforced microenvironment was considered, the heterogeneity of the fibrin gel substrate must also be acknowledged. As a common hydrogel system used in tissue engineering applications, the multiscale mechanical heterogeneity of fibrin gels has been well documented^51,52^. While we performed mechanical testing of overall fibrin gel formulations, heterogeneity is still present within the polymerization and crosslinking of the microscale fibrin microenvironment, which we have shown can influence the extent by which cells can respond. Second, although our clustering approach was successful in identifying responsive populations of MSCs used in our model system, the heterogeneity of cells must still be acknowledged. MSCs used in this study have not been definitively characterized, although it is reasonable to assume that only stromal cells adhered to the plastic dish during cell expansion.

## Conclusions

Utilizing a fiber-reinforced microenvironment system, we generated a large heterogeneous dataset of cells and utilized PCA and clustering to better reveal trends of cell response. Harnessing these principles, we showed that MSCs pattern the cell-fiber interface differentially according to their level of response, influencing how fiber-reinforced tissue engineering may need to optimize design at the microscale. In addition, we demonstrated that gel stiffness and matrix remodeling have a significant influence on how and where a cell responds, respectively. Overall, this research has several implications in aligned tissue engineering and can inform fiber-reinforced technologies built from the bottom up, utilizing microscale patterns of cell response to optimize design for organized and aligned tissue architecture.

## Acknowledgements

The authors thank the Department of Orthopaedics at Emory University and the Atlanta VA Medical Center for their support.

## Author Contributions

S.A.P., S.L.V., and J.M.P. designed experiments. S.A.P., M.H., H.S., and G.E.M. performed or assisted with biomaterial, cell culture, and imaging experiments. S.A.P., J.L.R., S.L.V., and J.M.P. interpreted data and results. All authors contributed to writing and editing the manuscript.

