## Supplemental Figures for "Revealing Early Spatial Patterns of Cellular Responsivity in Fiber-Reinforced Microenvironments"

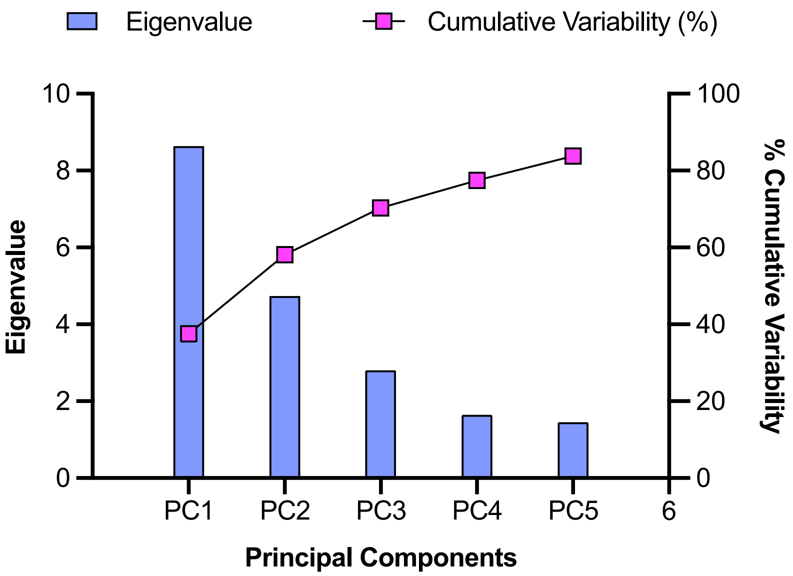


***Figure S1. Scree plot for 5 principal components***

*Eigenvalues and cumulative variability (%) for each of five principal components used for clustering following principal component analysis.*

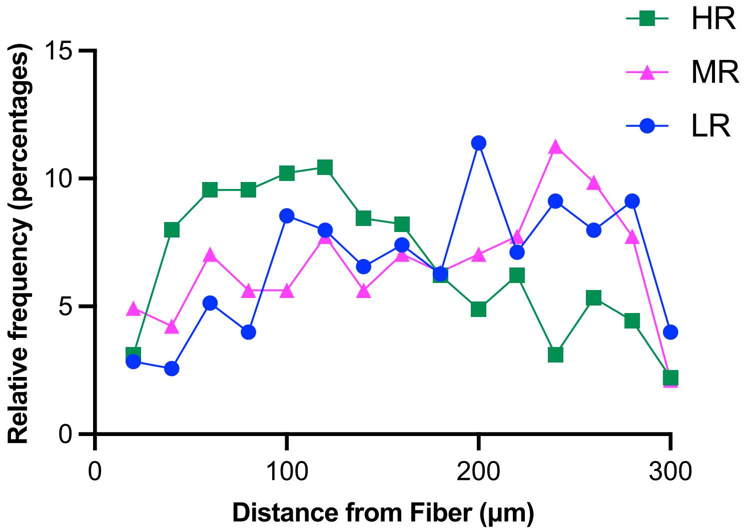


***Figure S2. Cells pattern area around fiber differentially based on cluster.***
*Frequency distribution histogram of cells in each cluster based on their distance from the fiber, organized into 20µm bins.*

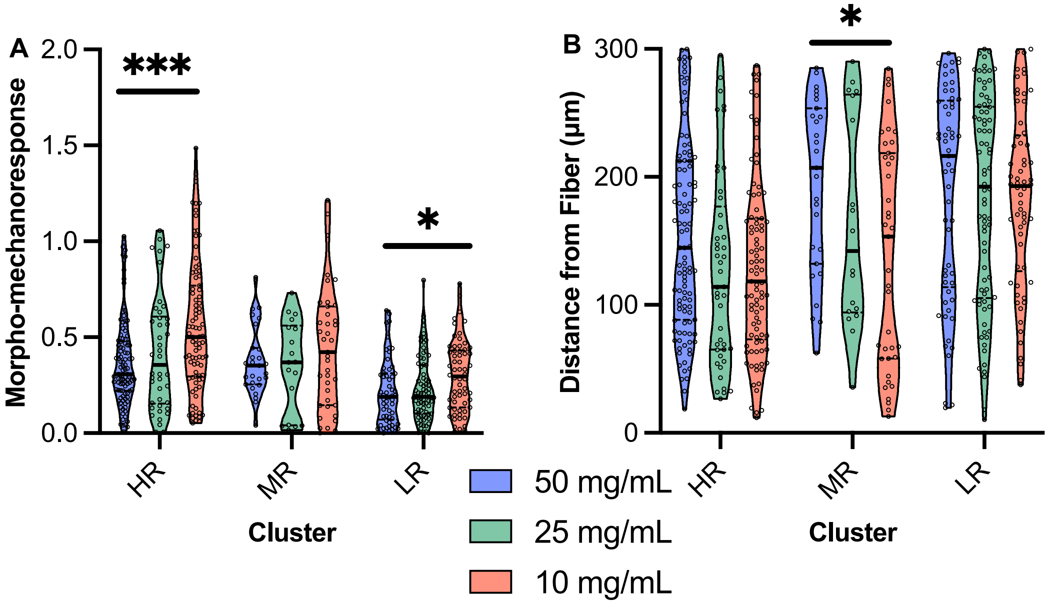


***Figure S3. Stiffness of hydrogel substrate variably influences cell morpho-mechanoresponse and spatial response of certain cell types in fiber-reinforced microenvironment.***
*[A] Morpho-mechanoresponse and [B] Distance from fiber of cells in fibrin gels of varying fibrinogen concentration, separated by cluster. *, *** represent p<0.05, 0.001, respectively.*

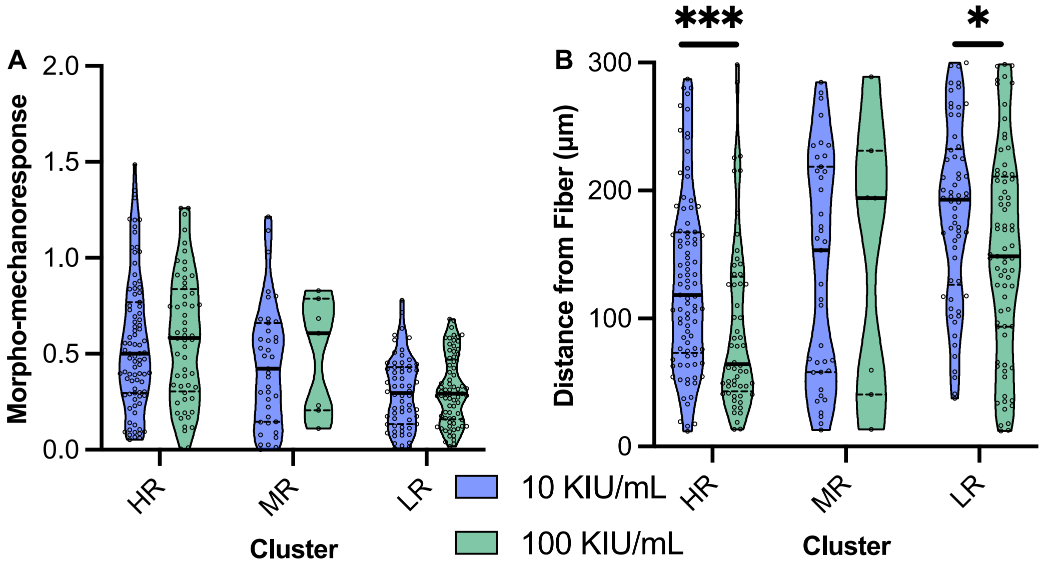


***Figure S4. Fibrin remodeling capacity during influences spatial response of certain cell types in fiber-reinforced micro-environment, but not individual cell mechanoresponse.*** *[A] Morpho-mechanoresponse and [B] Distance from fiber of cells in in fibrin gels supplemented with 10 and 100 KIU/ml concentrations of aprotinin, separated by cell cluster. *, *** represent p<0.05, 0.001, respectively.*
